# IL-4 and TGF-β Regulate Inflammatory cytokines and Cellular Infiltration in the Lung in Mouse-adapted SARS-CoV-2 Infection

**DOI:** 10.1101/2025.05.13.653138

**Authors:** Solomon Taye Sima, Lucinda Puebla-Clark, Maria Gonzalez-Orozco, Mark Joseph Endrino, Thomas R. Shelite, Hsiang-chi Tseng, Yazmin B. Martinez-Martinez, Matthew B. Huante, Hannah G. Federman, Komi Gbedande, Vineet D. Menachery, Mark C. Siracusa, Mark A. Endsley, Sara M. Dann, Janice J. Endsley, Ricardo Rajsbaum, Robin Stephens

**Affiliations:** Center for Immunity and Inflammation, Rutgers Health New Jersey Medical School, Newark, NJ, 07103, USA; Department of Internal Medicine, Division of Infectious Diseases, University of Texas Medical Branch, Galveston, TX 77555, USA; Department of Microbiology & Immunology, University of Texas Medical Branch, Galveston, TX 77555, USA; Department of Pathology, University of Texas Medical Branch, Galveston, TX 77555, USA; Center for Virus-Host-Innate Immunity and Department of Medicine; Department of Pharmacology, Physiology and Neuroscience, Rutgers Health New Jersey Medical School, Newark, NJ, 07301, USA

**Author notes:** These two authors made an equal contribution (Solomon Taye Sima, Lucinda Puebla-Clark).

**Keywords:** Cytokines, IL-4, TGF-β, Immunoregulation, SARS-CoV-2

## Abstract

The pathology of severe COVID-19 is due to a hyperinflammatory immune response persisting after viral clearance. To understand how the immune response to SARS-CoV-2 is regulated to avoid severe COVID-19, we tested relevant immunoregulatory cytokines. TGF-β, IL-10 and IL-4 were neutralized upon infection with mouse-adapted SARS-CoV-2 (CMA3p20), a model of mild disease; and lung inflammation was quantified by histology and flow cytometry at early and late time points. Mild weight loss, and lung inflammation including consolidation and alveolar thickening were evident 3 days post-infection (dpi) and inflammation persisted to 7 dpi. Coinciding with early monocytic infiltrates, CCL2 and granulocyte-colony stimulating factor (G-CSF) were transiently produced 3 dpi, while IL-12 and CCL5 persisted to 7 dpi, modeling viral and inflammatory phases of disease. Neutralization of TGF-β, but not IL-10 or IL-4, significantly increased lung inflammatory monocytes and elevated serum but not lung IL-6. Neutralization of IL-4 prolonged weight loss and increased early perivascular infiltration without changing viral titer. Anti-IL-4 reduced expression of *Arg1*, a gene associated with alternative activation of macrophages. Neutralizing TGF-β and IL-4 had differential effects on pathology after virus control. Lung perivascular infiltration was reduced 7 dpi by neutralization of IL-4 or TGF-β, and peri-airway inflammation was affected by anti-TGF-β, while alveolar infiltrates were not affected by either. Anti-IL-4 prolonged IL-12 to 7 dpi along with reduced IL-10 in lungs. Overall, the immunoregulatory cytokines TGF-β and IL-4 dampen initial inflammation in this maSARS-CoV-2 infection, suggesting that promotion of immunoregulation could help patients in early stages of disease.

**Visual Abstract:** 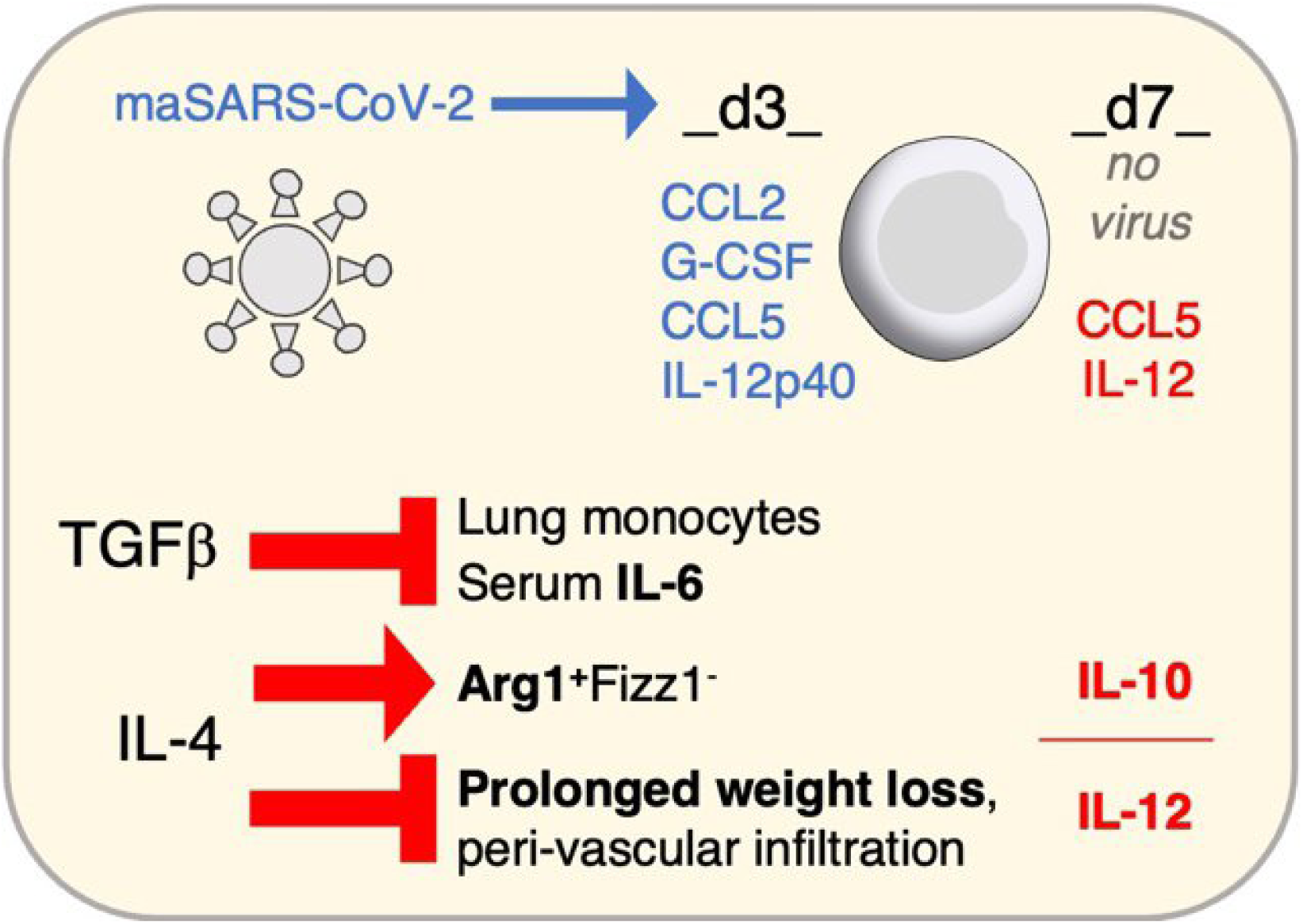

## Introduction

Infection with severe acute respiratory syndrome-coronavirus-2 (SARS-CoV-2) causes coronavirus disease 2019 (COVID-19), which has caused over 7 million deaths worldwide in four years ^1^. The immune response to SARS-CoV-2 infection in has two phases: a viral control phase and a hyperinflammatory phase, which can cause severe disease if not well regulated. The severity of COVID-19 is dependent on SARS-CoV-2 viral load, variant tropism and the intensity and duration of the host immune response. The virus initially causes tissue damage in the upper respiratory tract, which can progress into the lungs ^2, 3^. After an illness lasting 1-2 weeks, most patients resolve the infection; however, up to 40% of admitted patients require intensive care due to severe pulmonary complications, which can lead to death ^4^. During this period after acute virus infection, high levels of systemic inflammatory markers, such as interleukin (IL)-6 and Complement-reactive protein, correlate with severity and prolonged hospital stays ^3, 5^. While there is evidence of persistent virus, the importance of prolonged inflammation in severe disease is supported by the first therapeutic successes, namely immunosuppressants, including corticosteroids and the Jak1/2 inhibitor baricitinib, which are still therapeutic after hospital admission ^6, 7^. This prolonged inflammatory phase of disease has unique aspects compared to influenza or respiratory syncytial virus suggesting unique mechanisms of pathogenesis that remain poorly understood at the immunoregulatory level ^8^. Here we are using a mouse adapted virus first described by Muruato, *et al* ^9^ which peaks at day 2 of infection, is live in the lungs at day 4, but is undetectable by plaque assay or PCR by day 7. This virus causes mild disease suggesting that sufficient immunoregulation during the peak of infection regulates the potential for persisting immune-mediated damage ^10^.

The immune response to SARS-CoV-2 infection begins with a limited interferon (IFN)-I response due to successful viral antagonism ^11–13^ in the presence of an early burst of TNF and IL-1. This systemic inflammatory response coincides with lung infiltration consisting of neutrophils, monocytes and dendritic cells, as occurs in humans ^14^. Severe disease corresponds to elevated production of inflammatory cytokines from vascular endothelium, respiratory epithelium and immune cells ^3, 15^. Serum IL-6 is reliably associated with disease severity ^16^; though it is unclear what regulatory cytokines modulate IL-6 in vivo ^17, 18^. The correlation of IL-6 with disease severity led to early clinical trials studying potential COVID-19 therapies; however, anti-IL-6 antibodies were not successful as therapy once patients were admitted, suggesting that the role of IL-6 in monocytic inflammation occurs before patients arrive at the hospital ^19–21^. Novel populations of monocytes are recruited to the lungs in severe COVID-19 ^22^. CCL2 and its receptor CCR2 are essential for monocyte trafficking from the bone marrow under inflammatory conditions ^23^, while CCL5 is thought to locally recruit inflammatory monocytes to tissues through their expression of CCR5 ^24–26^. CCL5 and CCL2 are implicated in COVID-19 pathogenesis and monocyte-derived cells play a protective role against mouse-adapted SARS-CoV-2 (maSARS-CoV-2) ^27, 28^.

As it is so crucial to understand how to limit inflammation in the response to SARS-CoV-2, we targeted immunoregulatory cytokines TGF-β, IL-10 and IL-4 to assess their roles in pathology in maSARS-CoV-2 infection. While this is not a model of severe disease, there is evidence of monocyte recruitment and lung inflammation ^14, 23^. TGF-β and IL-10 are the most well understood immunoregulatory cytokines, known to be essential in very inflammatory conditions such as extracellular parasite infections and inflammatory bowel disease ^29^. We also tested IL-4 due to evidence that Th2-type cytokines and alternatively activated macrophages are involved in human and mouse SARS-CoV-1 and -2 infections ^30–37^.

Evidence of Th2-type immune response to SARS-CoV-2 includes evidence that Eosinophilia and IL-13 correlate with severity in COVID-19 ^31^, and that helminth infection reduces prevalence of COVID-19 ^32^. In addition, IL-5 and IL-13 are expressed in patients and mice infected with SARS-CoV-2 ^33^. However, IL-5 and IL-13 are considered Th2 effector cytokines, and neither has been shown to regulate IFN-I or Th1-type inflammation ^34^. On the other hand, the signature Th2 cytokine IL-4 can inhibit the synthesis of the proinflammatory mediators TNF-α, IL-1, IL-6, IL-12, PGE_2_, and the chemokines IL-8 and MIP-1 ^35^, as well as the differentiation of type-1 T helper lymphocytes ^36^. As Th2 cytokines are also involved in wound repair ^37^, we are interested in the role of these cytokines in the phase of disease that determines the early severity and then the later phase of disease where lung tissue homeostasis is restored. Therefore, in this study we studied the role of immunoregulatory cytokines during the peak of infection, as well as after virus is largely cleared.

The data indicate regulatory roles for IL-4 and TGF-β infection with mouse adapted SARS-CoV-2. Neutralization of TGF-β throughout infection dramatically increased cellular infiltration in all lung spaces at 3 dpi and increased systemic IL-6 and IL-22. Inhibiting IL-4 also increased early cell infiltration, and prolonged weight loss and increased late IL-12p40 and CCL5. We showed upregulation of *Arg1* in lung tissue, suggesting alternative activation of macrophages in maSARS-CoV-2 infection, as previously shown in SARS-CoV infected lungs ^30^. *Arg1* expression was dependent on IL-4, which was present in the lung at low levels before infection. Neutralizing either TGF-β or IL-4 changed the pattern of inflammation seen in the lungs at 7 dpi, with IL-4 limiting infiltration specifically at peri-vascular locations at the later time point. Regulatory mechanisms will need to be further investigated to understand their potential to contribute to limiting the severity of COVID-19.

## Materials and Methods

### Mice and in vivo antibody injection

C57BL/6J mice were purchased from the Jackson Laboratory, maintained, and aged with ad libitum access to food and water, under specific-pathogen-free conditions in the Animal Resource Center facility at University of Texas Medical Branch (UTMB). Male mice 38 to 41-weeks old were infected under animal biosafety level 3 (ABSL-3) conditions at UTMB or Rutgers NJMS in accordance with institutional biosafety approvals. Mice were treated intraperitoneally (i.p.) with either 400μg anti-mouse IL-4 (Clone 11B11; BP0045, BioXcel, Lebanon, NH), 200μg anti-mouse TGF-β (1D11.16.8; BE0057, BioXcel), or mixed isotype controls (400μg rat IgG1 for IL-4, TNP6A7; BE0290, and 200μg mouse IgG1 for TGF-β, MOPC-21; BE0083, BioXcel)), every other day starting the day of infection (days 0, 2 and 4) under anesthesia. Isotype controls were also given to mock-infected animals. All animal experiments were carried out in accordance with the Institutional Animal Care and Use Committee (IACUC) guidelines and have been approved by both the UTMB and Rutgers IACUC.

### Virus infection

Mouse-adapted SARS-CoV-2 (CMA3p20) was grown in Vero E6 cells, as described by Muruato, *et al* ^9^. Mice were anesthetized with 5% isoflurane and infected intranasally with 50μL of media containing 1x10^6^ plaque forming units (PFU) of SARS-CoV-2 (CMA3p20). Mice in the mock group were given only phosphate-buffered saline (PBS) intranasally. Health checks and body weight were recorded daily. Lungs and serum were collected for downstream analysis.

### Single-cell suspension and flow cytometry

Lungs were collected in RPMI (11875-093) 10% v/v FBS (S12450H) and 1% v/v penicillin-streptomycin (both from ThermoFisher), and washed in PBS, followed by enzymatic digestion with collagenase D and DNase I (both from Roche, Mannheim, Germany) in serum-free RPMI for 30 min in a humidified 5% CO_2_ incubator at 37°C. After incubation, FBS was added to stop the enzymatic activity. The single-cell suspension was collected by passing the cells through 70μm strainer; and red blood cells were lysed with RBC lysis buffer (Life Technologies, Carlsbad, CA). 1x10^6^ cells were stained using the following antibodies: Anti-CD4-BV421 (clone RM4-5), CD25-BV785 (PC61.5), MHCII-BV605 (M5/114,15.2), CD163-BV421 (S15049I), CD206-PE (MR6F3), CD45-PerCP-Cy5.5 (30-F11), CD11c-PE (N418), Siglec-F-BV421 (S17007L), Ly6C-BV650 (HK1.4), CD11b-BV780 (M1/70). Live-Dead Fixable blue stain (Life Technologies, OR) was used to exclude dead cells. After staining, samples were inactivated and fixed with 4% ultrapure formaldehyde (18814, PolySciences, PA) for 24hrs, and then the cells were resuspended in 2% ultrapure formaldehyde for another 24hrs and acquired on the LSR II Fortessa Custom 2/5/3/5/2 (Becton Dickinson, CA) and analyzed with FlowJo, version 10.8.1 (Becton Dickinson, OR), similar to ref ^10^.

### Virus quantification

Infectious virus in lung tissue was quantified by plaque assay. Tissue was collected in 1mL of DMEM (11965092, ThermoFisher, Waltham, MA) 10% FBS. Tissue samples were weighed and homogenized, and then quickly frozen. Six-fold serial dilutions of homogenates were prepared in duplicate and used to inoculate 4x10^5^ Vero E6 cells grown in a 6-well tissue culture plate containing 1mL DMEM 10% FBS. Following 1hr incubation at 37°C 5% CO_2_, the cells were overlayed with 1.6% Tragacanth solution (G1128, Sigma, St Louis, MO) supplemented with 2% FBS and 1X penicillin-streptomycin antibiotics and incubated for 2 days. Cells were then fixed in 10% formalin and plaques were visualized by staining with 1% crystal violet (V5265, Sigma) diluted in 10% ethanol (1070172511, Sigma). Viral infectivity titers were expressed as PFU/mL in Vero E6 cells and were calculated according to the Behrens-Karber method.

### Real-time quantitative PCR

Total RNA was isolated from lung using Direct-zol RNA MiniPrep (Zymo Research, Irvine, CA) according to the manufacturer’s protocol. Reverse transcription was conducted using high-capacity cDNA reverse transcription kit (Applied Biosystems, Foster city, CA) according to the manufacturer’s protocol. Primers and SYBRGreen mix were purchased from Bio-Rad (Hercules, CA) and some primers were from Origene (Rockville, MD). DNA was amplified with QuantStudio 6 Flex real-time PCR System (Applied Biosystems, Foster City, CA).

### Cytokine Bead Assay

Luminex Intelliflex using MAGPIX beads was run according to the manufacturer’s protocol. High sensitivity (IFN-γ, IL-2, IL-4, IL-6 and TNF-α) and the custom cytokine (IL-1α, IL-10, IL-12-p40, IL-13, IL-18, IL-22, IL-25, IL-27, IFN-β, CCL2, CCL5, CXCL2, TSLP and G-CSF) panel kits (Bender MedSystems, Vienna, Austria) were processed according to manufacturer’s instructions. The analysis was performed using the Luminex Intelliflex analysis software (Thermo Fisher, Waltham, MA).

### Histology

The right inferior lobe of the lung was fixed in 10% neutral buffered formalin (HT501128, Sigma, MI) for 7 days. Tissues were cut, paraffin blocks were made, and H&E staining performed by the Anatomic Pathology Laboratory of the Pathology Department of University of Texas Medical Branch. The pathology scoring was conducted in a blinded manner. Pathology score was determined using an observational scale developed from the sections. Interstitial Thickening was scored 1 if yes plus 1 if < 20% of lung affected, 2 if 25-50% and 3 if 60-75%. Peribronchiole infiltrates were scored 1 if yes plus scored 1 if yes plus 1 if 1-2 foci, 2 if 2-3 and 3 if 3 or more. Perivascular infiltrates 1 if yes plus scored 1 if 1-2 foci, 2 if 2-3 and 3 if 3 or more. All infiltrates were scored 1 if consolidation present, or 1 if 1-2 areas and 2 if 3 or more areas present. Additionally, one was added for each of fibrolytic tissue, Edema or debris or alveolar fluid, if present. The maximum score observed was 19. Peri-vascular, peri-airway and nuclei counting for thickness were done manually in a blinded manner.

### Statistical analysis

FlowJo 10.8.1 and GraphPad Prism version 9.1.1 (GraphPad Software) were used to perform data and statistical analysis. Statistical analysis was performed by one-way ANOVA followed by Student’s t test. Intergroup comparisons were performed using nonparametric analysis. All data are expressed as the mean and standard error of the mean (SEM). The statistical details of the experiments are provided in the respective figure legends. A *p* value at < 0.05 was considered statistically significant. \**p*=0.01-0.05, \*\**p*=0.01-0.001, \*\*\**p*=0.001-0.0001, \*\*\*\**p*<0.0001.

## Results

Viral ORF1ab RNA was detected in the lung by real-time qPCR on 2, but not 7 dpi (**Fig. 1A**), and has been reported to remain detectable by PCR and plaque assay through day 4 ^38^. To characterize the immune response to this mouse-adapted virus infection, cytokines and chemokines were measured in lung homogenate and serum of infected mice by multiplex bead-based assay on 3 and 7 dpi. Cytokines and chemokines related to differentiation and recruitment of monocytes present in the serum were IL-6, G-CSF and the chemokines CCL2 (MCP1) and CCL5 (RANTES) ^39–41^. The infection induced G-CSF and CCL2 on 3 dpi, while CCL5 increased compared to mock-infected by 7 dpi (**Fig. 1B**). IL-10 was not increased above background on 3 dpi (**Fig. 1C**), while IL-12 p40 was increased by infection and remained elevated in isotype-treated mice until 7 dpi (**Fig. 1D**). IL-33, an epithelial and endothelial cell damage alarmin, was induced by the infection on 3 dpi, but returned to baseline by 7 dpi (**Fig. 1E**). Histopathological evaluation of the lungs on 3 dpi showed that mice infected with maSARS-CoV-2, that also received isotype control antibody, had more leukocytes infiltration into the alveoli, and around airways and vessels than in mock group animals (**Fig. 2A**). There was also alveolar thickening and consolidation seen in the infected lungs on 3 dpi. The infiltration was determined microscopically to be highly monocytic, consistent with findings in humans ^22^. The pathology score shows that both isotype and anti-IL-4 have more severe lung damage than mock group on 3 dpi (**Fig. 2B**). No differences in lung pathology were observed between anti-IL-10 and isotype control treated animals (not shown); therefore, not all of the samples from this group were processed for all assays.

**Figure 1.**
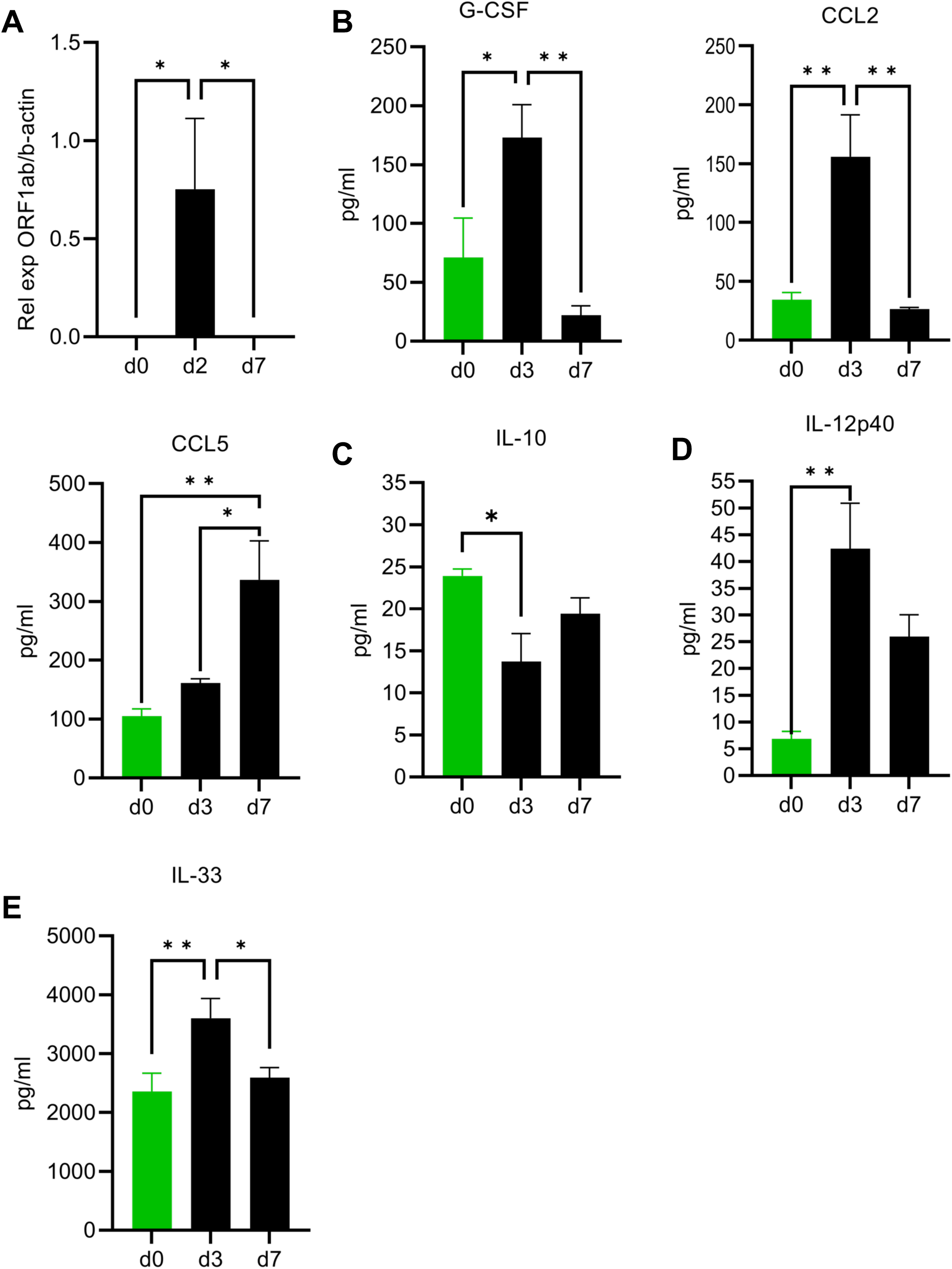
Weight loss and cytokine production in aged B6 mice infected with maSARS-CoV-2. C57BL/6 mice were infected at ten months of age with maSARS-CoV-2, or mock and treated with isotype control antibody on alternate days and weighed daily. Lung homogenate was taken for measuring cytokines on day 3 p.i. Graphs show; **A)** viral load by qPCR; **B)** monocyte-related cytokines and chemokines G-CSF, CCL2, CCL5; **C)** IL-10; **D)** IL-12p40; **E)** IL-33. Results are representative of at least two experiments. Statistical analysis was performed by one-way ANOVA followed by Student’s t test. Data were presented as mean ± SEM for groups of five mice in mock and isotype on d0, d3 and d7 p.i. A *p* value at <0.05 was considered statistically significant.

**Figure 2.**
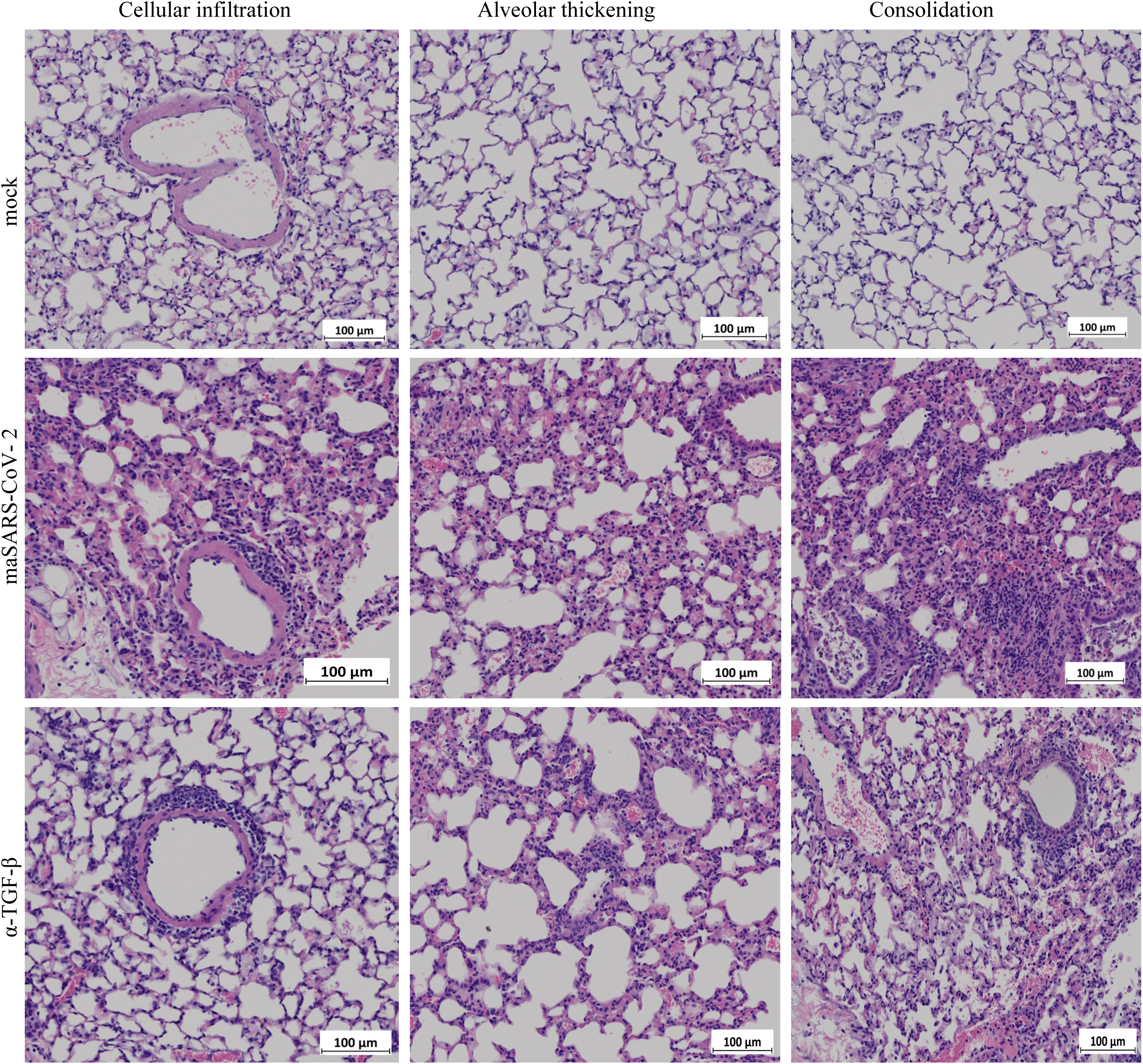
Lung pathology of maSARS-CoV-2 infected mice. C57BL/6 mice were infected at ten months of age with maSARS-CoV-2, or mock and treated with isotype control or TGF-β neutralizing antibodies on alternate days and weighed daily. H&E of lungs of mock (top), isotype control (middle) and TGF-β (bottom). Results are representative of at least two experiments. Cellular infiltration, alveolar thickening and consolidation were identified, and representative fields are shown. Scale bars represent 100μm.

### TGF-β regulates cellular infiltration into the lung

To determine the effects of TGF-β on acute and persistent inflammation in maSARS-CoV-2 infection, we studied the pathology of the lungs after neutralization of TGF-β. H&E staining of lungs from infected mice treated with anti-TGF-β showed an increase in leukocytes in the alveolar walls 3 dpi, with alveolar thickening and some consolidation, or filling of some alveolar airspaces (**Fig. 2A**). Thickening was consistent throughout the infected lung section, while the fraction of the lung consolidated in each animal was variable.

To determine the cellular composition of the infiltrate, flow cytometry was performed on lungs from infected mice 3 dpi. Neutralization of TGF-β increased infiltration of CD45^+^ leukocytes in the lungs. Three populations of CD11b^+^ cells were identified using side scatter characteristics (**Fig. 3A**). The least granular (SSC^lo^) CD11b^+^ cells contained the Ly6C^hi^ (CD11c^-^) inflammatory monocytes, while the most granular, or vacuolated cells (SSC^hi^) contained the CD11c^+/int^ alveolar macrophages. Quantifying these results, we found that anti-TGF-β treatment increased infiltration of CD45^+^ leukocytes, with a specific increase in inflammatory monocytes (CD11b^+^SSC^lo^Ly6C^+^) (**Fig. 3B**). The SSC^int^ population contained Ly6C^int^ cells and may be regulatory, as it decreased in the absence of IL-4. SSC^int^CD11b^+^ cells were not further identified, and Ly6G was not included in the panel to identify neutrophils as they were rarely identified in H&E tissue sections. The number of alveolar macrophages (CD11b^+^SSC^hi^) was decreased in the anti-TGF-β group.

**Figure 3.**
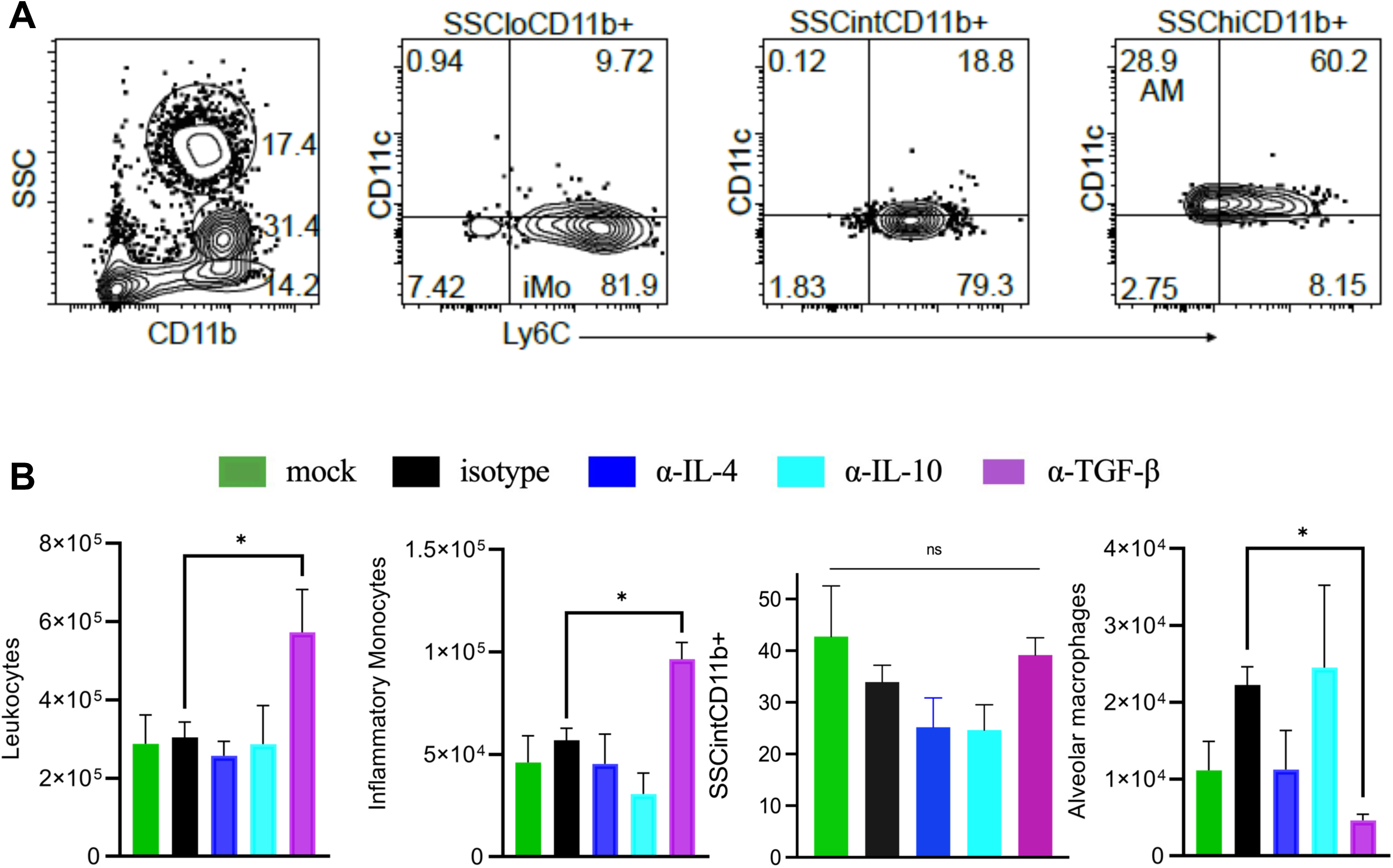
TGF-β regulates cellular infiltration in the lung. C57BL/6 mice were infected with maSARS-CoV-2 and treated with isotype or cytokine neutralizing antibodies on alternate days and weighed daily. **A)** Flow cytometry plots showing gating on CD11b^+^ live cells with three levels of side scatter in the lungs on day 3 p.i and Ly6C and CD11c staining in those gates for identification; **B)** quantification of infiltrating leukocytes (CD45^+^), inflammatory monocytes (CD11b^+^SSC^lo^), undefined monocytic cells (CD11b^+^SSC^int^) and alveolar macrophages (CD11b^+^SSC^hi^). Results are representative of at least two experiments. Statistical analysis was performed by one-way ANOVA followed by Student’s t test. Data were presented as mean ± SEM for groups of five mice per group. A *p* value at <0.05 was considered statistically significant.

TGF-β was detected in the lungs but was not induced by infection at 3 or 7 dpi (**Fig. 4A**). Neutralization of TGF-β did not change live virus in the lung tissue significantly compared to the isotype treated-infected group, though there was a significant trend downward in multi-group comparison (**Fig. 4B**). IFN-γ was induced 3 dpi, but did not vary between infected groups (**Fig. 4C**). As IL-6 varies considerably between patients, correlates with disease severity ^42^, and also plays a role in monocyte differentiation ^20, 43^; we measured this critical cytokine in lung homogenate and serum by bead-based ELISA assay. IL-6 was induced in the lung upon infection at 3 dpi, but did not vary significantly among groups (**Fig. 4D**). Although we did not see an increase in serum IL-6 between mock infected and infected isotype control animals in these experiments, IL-6 was significantly higher in the serum of infected mice treated with anti-TGF-β than in the isotype group (**Fig. 4E**). IL-12 p40 was increased in the lung on 3 dpi (**Fig. 4F**). This data supports a significant role for TGF-β in the regulation of both acute lung infiltration and systemic cytokine storm.

**Figure 4.**
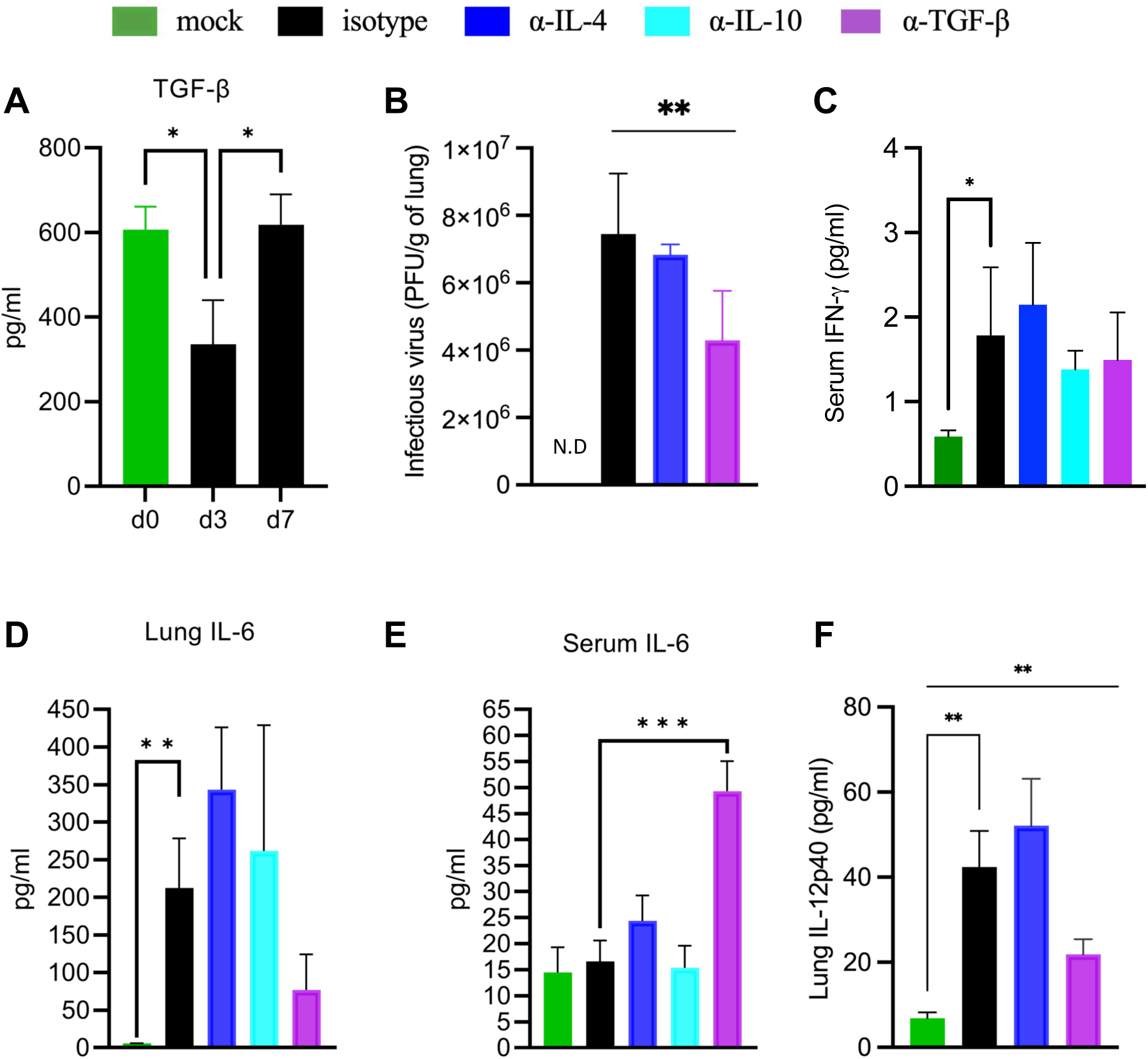
TGF-β regulates IL-6 and modulates viral clearance. Aged C57BL/6 mice were infected with maSARS-CoV-2 and treated with isotype or cytokine neutralizing antibodies on alternate days and weighed daily. On day 3 p.i., the lung homogenate and serum were tested for virus and cytokines. Cytokine bead assay shows cytokine levels of **A)** TGF-β; **B)** viral quantification; **C)** IFN-γ in serum **D)** IL-6 in lung homogenate and **E)** in serum; **F)** IL-12p40 in lung homogenate. Results are representative of at least two experiments. Statistical analysis was performed by one-way ANOVA followed by Student’s t test. Data were presented as mean ± SEM for groups of five mice. A *p* value at <0.05 was considered statistically significant. n.d. not detected.

### Neutralizing IL-4 prolongs pathology

The effect of immunoregulatory cytokines on weight loss, which is a sign of clinical disease, was assessed. While infected, isotype-control treated mice showed significant weight loss that had a small peak 3 dpi. Neutralization of IL-4 led to weight loss that was prolonged compared to isotype-control, with significantly more weight loss at 4 dpi than isotype-control (**Fig. 5A**). Virus quantification by plaque assay was similar in anti-IL-4 treated mice to the isotype control group by qPCR (**Fig. 5B**). IL-4 was found to be expressed even in the lungs of mock-infected mice when tested with a high-sensitivity bead-based assay, suggesting low homeostatic levels (**Fig. 5C**). IL-13 was also detected in the isotype control group on both 3 and 7 dpi and was not higher than the mock-infected group in these experiments (**Fig. 5D**).

**Figure 5.**
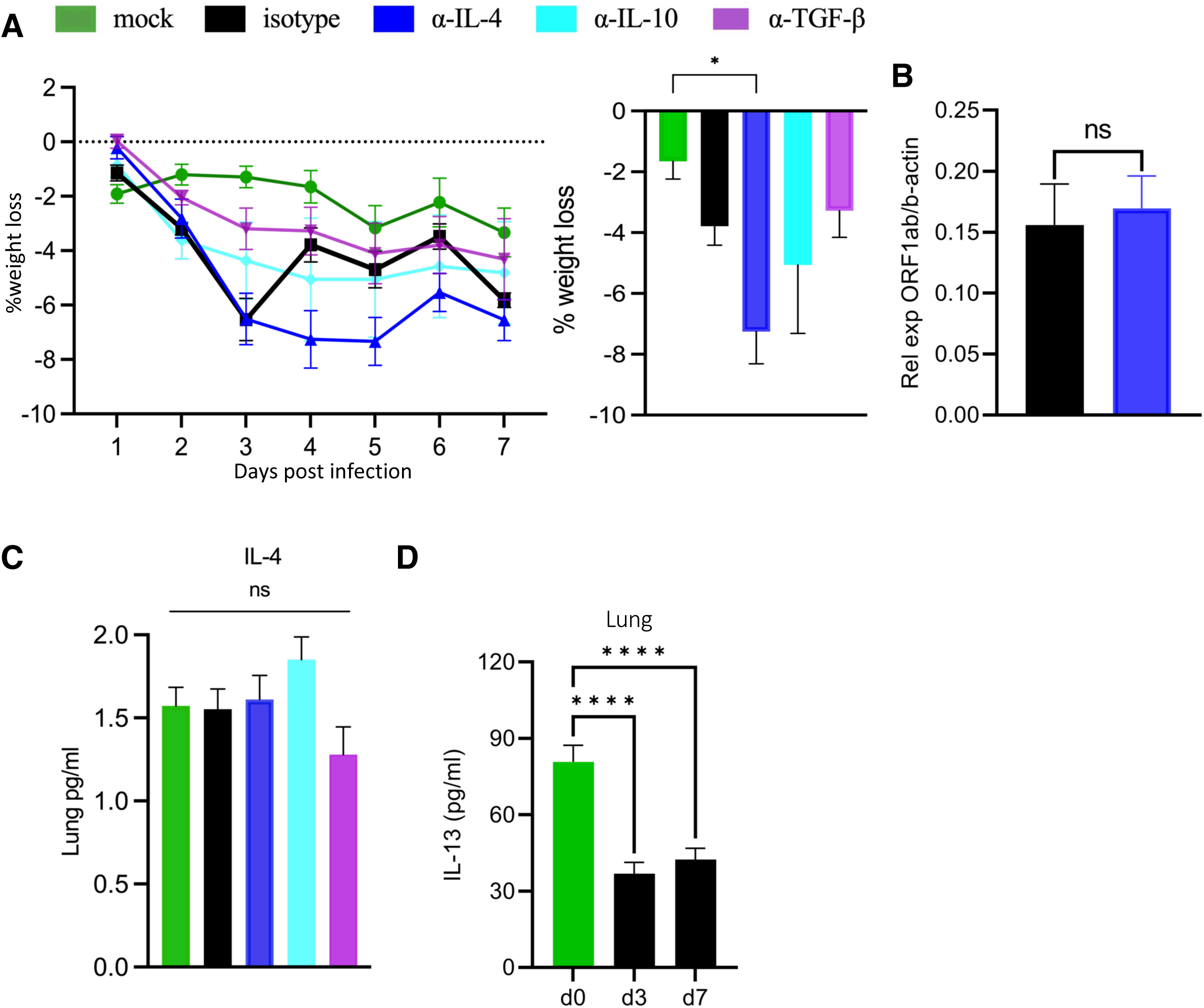
Neutralization of IL-4 prolongs weight loss without change in viremia. Aged C57BL/6 mice were infected with maSARS-CoV-2 and treated with isotype or cytokine neutralizing antibodies on alternate days and weighed daily. **A)** Percent weight loss over the course of infection, and statistics on day 4 p.i; **B)** quantification of virus on day 3 p.i by relative levels of *ORF1ab* by qPCR; Cytokine levels in lung homogenate **C)** IL-4 on day 3 p.i. and **D)** IL-13 are shown in bar graphs. Results are representative of at least two experiments. Statistical analysis was performed by one-way ANOVA followed by Student’s t test. Data were presented as mean ± SEM for groups of five mice per group. A *p* value at <0.05 was considered statistically significant.

Neutralizing IL-4 has an effect on lung inflammation. Anti-IL-4-treated mice showed increased infiltration in perivascular spaces and around airways 3 dpi as shown in **Figure 6A**. Alveolar thickening, related to alveolar infiltration, and consolidation were similar to isotype control. All of these observations are quantified as an overall pathology score in **Figure 6B**. Anti-IL-4, but not anti-TGF-β showed these effects on peri-ariway and -vascular infiltration as quantifed in **Figure 6C**. There was no significant difference in alveolar cellularity between isotype and anti-IL-4, or anti-TGF-β-treated groups 3 dpi (**Fig. 6D**). Collectively, these data support a role for IL-4 in regulating lung pathology in this infection. As IL-4 can drive alternatively activated macrophage differentiation in infection ^31^, we tested for genes correlated with this phenotype in RNA from infected lungs on day 3 p.i. Arginase 1 (*Arg1*), an AAM effector gene, was expressed in infected lungs, and significantly more in isotype than anti-IL-4-treated animals (**Fig. 6E**). However, expression of resistin-like molecule a1 (*Fizz1*), another AAM transcript, was not increased in infection 3 dpi (**Fig. 6F**).

**Figure 6.**
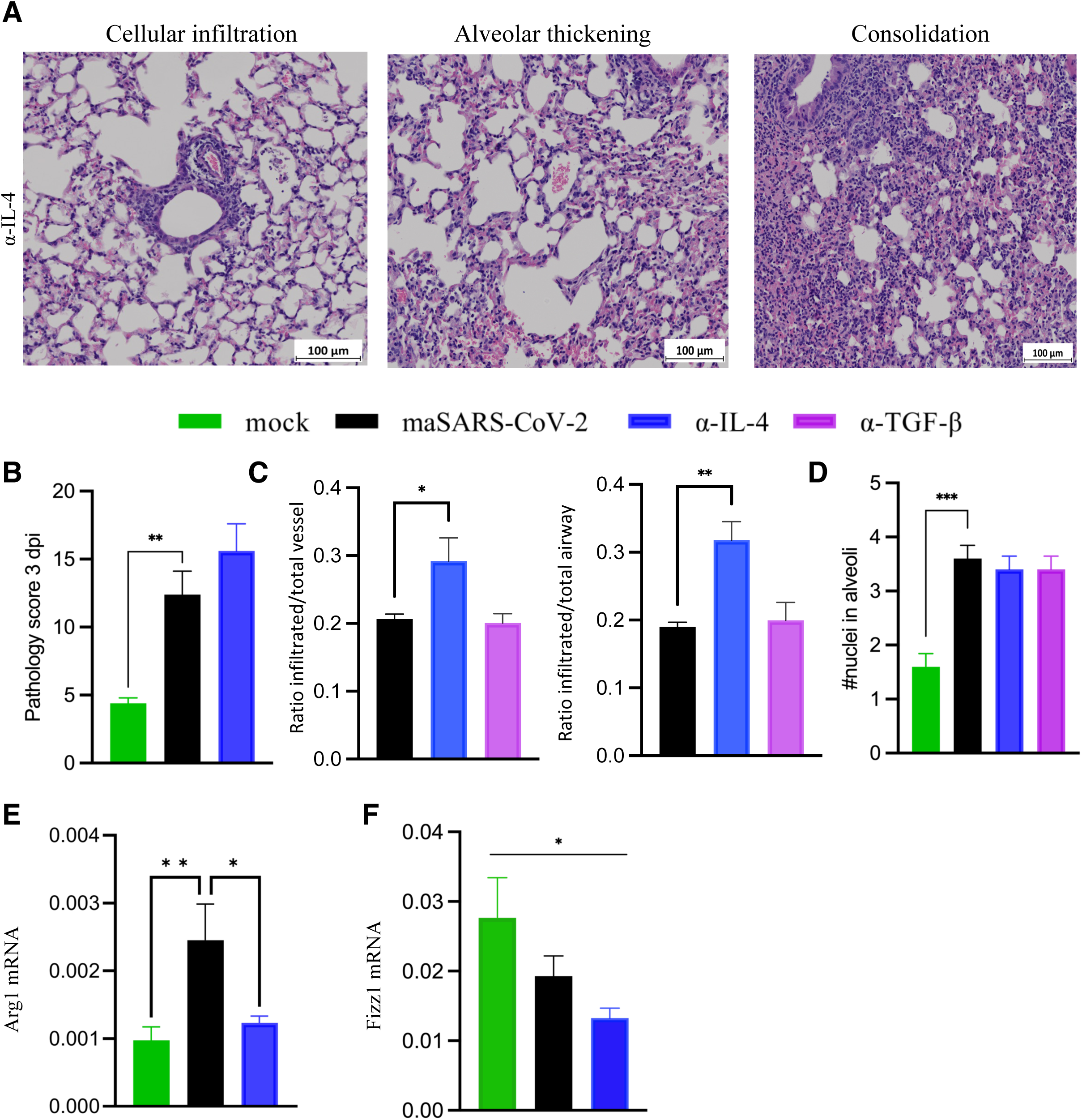
Neutralization of IL-4 leads to increased cellular infiltration on 3 dpi. Aged C57BL/6 mice were infected with maSARS-CoV-2 and treated with isotype or cytokine neutralizing antibodies on alternate days and weighed daily. **A)** H&E staining of lungs day 3 p.i. from mice treated with anti-IL-4. Results are representative of at least two experiments. Cellular infiltration, alveolar thickening and consolidation were identified, and representative fields are shown. Scale bars represent 100μm; **B)** Quantification of overall pathology score from day 3 p.i. **C)** Quantification of peri-vascular and peri-airway infiltration using ratio of inflamed to total; **D)** Alveolar infiltration represented by a score based on average number of nuclei in the thickness of the alveoli in each field. Statistical analysis was performed by one-way ANOVA followed by Student’s t test. Data were presented as mean ± SEM for groups of five mice. A *p* value at <0.05 was considered statistically significant.; **E)** *Arg1* and **F)** *Fizz1* expression in the lungs by real time PCR. Results are representative of at least two experiments. Statistical analysis was performed by one-way ANOVA followed by Student’s t test. Data were presented as mean ± SEM for groups of five mice. A *p* value at <0.05 was considered statistically significant.

As we had obeserved elevated inflammatory cytokines CCL5 and IL-12p40 7 dpi, after virus became undetectable, we evaluated lung histology on 7 dpi (**Fig. 7A**). All infected groups still had some signs of inflammation at this timepoint. Significant interstitial thickening with 60-75% of the lung affected was observed in most infected mice treated with isotype control (3/5). Peribronchial and perivascular infiltrates were present in all mice, with half of the mice having more than three foci of inflammation in the lobe section studied. Consolidation was present in all mice, and half of the mice exhibited consolidation in three or more areas. Half of the mice had detectable necrotic cells in areas of consolidation and one mouse had alveolar edema. However, treatment with anti-IL-4 and anti-TGF-β changed the intensity of infiltration as well as the spatial distribution of cells and damage. The histopathology and pathology score showed signs of recovery by 7 dpi in the lungs of the anti-IL4 treated group; However, the pathology score showed that the infected, isotype-treated group still had more lung damage than mock-infected animals 7 dpi (**Fig. 7B**). Perivascular infiltration was specifically reduced in mice given neutralizing antibodies to the immunoregulatory cytokines TGF-β and IL-4, compared with isotype-treated groups, while peri-airway infiltration was specifically reduced by anti-TGF-β at this late timepoint suggesting that they may inhibit resolution of pathology (**Fig. 7C**). Alveolar infiltration, the most likley feature to be associated with reduced lung function, remained observable at 7 dpi and was not significantly affected by neutralization of regulatory cytokines (**Fig. 7D**). These quantitative studies did not include lungs from the anti-IL-10 treated mice, which appeared grossly similar to isotype control. Combining observations at the two time points, it appeared that neutralizing IL-4 shifts the peak of infiltration earlier, at least around vessels, suggesting early regulation by this cytokine, which is supported by prolonged weight loss at 4 dpi separating the events in the pathological cascade into two mechanistic phases, as in humans.

**Figure 7.**
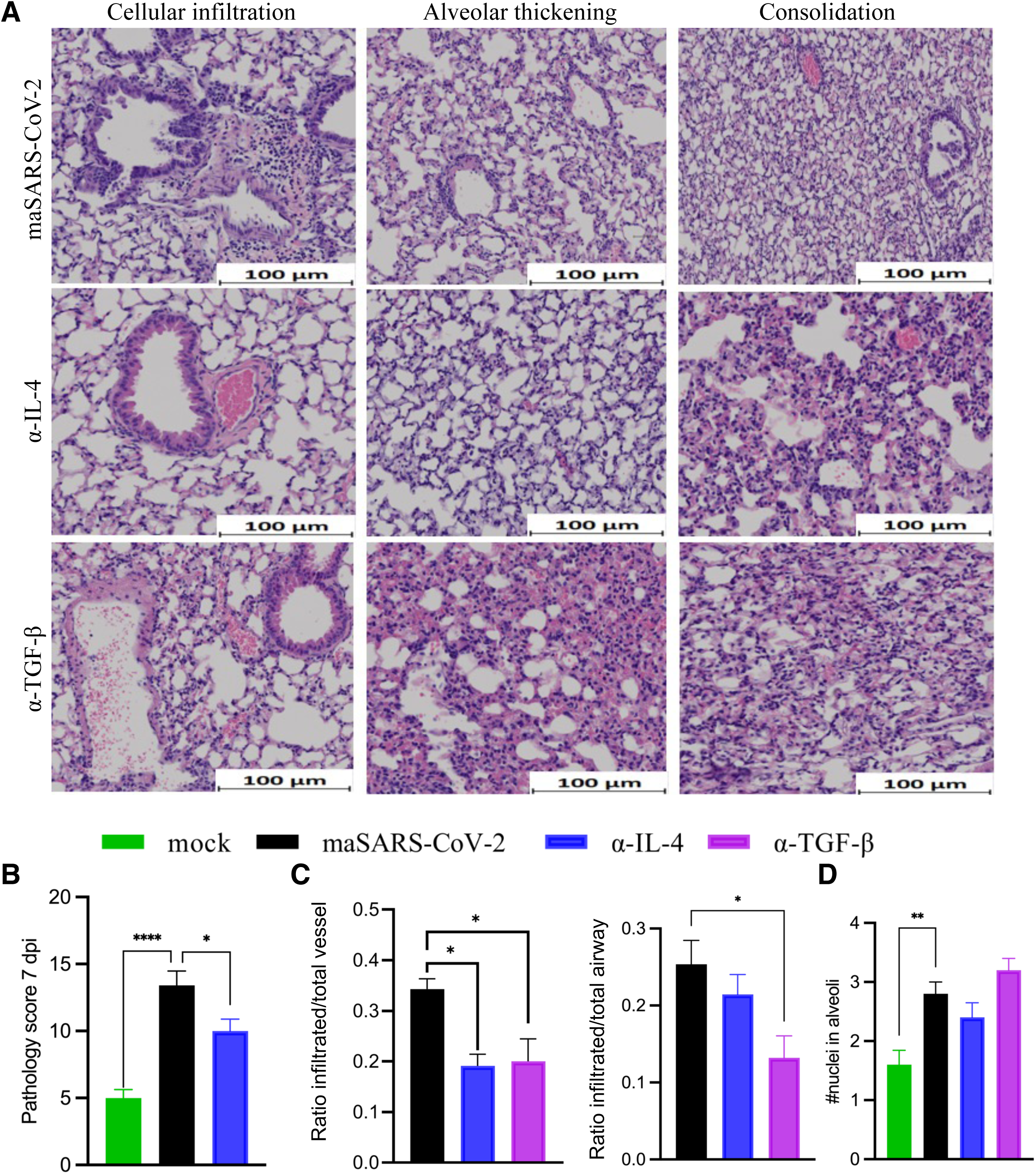
Lung pathology of maSARS-CoV-2 infected mice 7 dpi. C57BL/6 mice were infected at ten months of age with maSARS-CoV-2 and treated with isotype control or cytokine neutralizing antibodies on alternate days and weighed daily. **A)** H&E of lungs of isotype (top), IL-4 (middle) and TGF-β (bottom). Results are representative of at least two experiments. Cellular infiltration, alveolar thickening and consolidation were identified, and representative fields are shown. Scale bars represent 100μm; **B)** Quantification of overall pathology score 7 dpi; **C)** Quantification of peri-vascular and peri-airway infiltration using ratio of inflamed to total, and **D)** alveolar infiltration represented by a score based on average number of nuclei in the thickness of the alveoli in each field on 7 dpi. Statistical analysis was performed by one-way ANOVA followed by Student’s t test. Data were presented as mean + SEM for groups of five mice per group. A p value of <0.05 was considered statistically significant.

To test potential causes of persistent inflammation 7 dpi, we measured the monokines CCL5 (**Fig. 8A**), IL-12p40 (**Fig. 8B**) and IL-10 (**Fig. 8C**), as well as IFN-γ (**Fig. 8D**). Several appeared upregulated, but only IL-12 p40 reached significance in the lung of anti-IL-4 treated mice when compared to the isotype control infected group on 7 dpi. Strikingly, in the anti-IL-4-treated group, IL-10 was reduced compared to isotype on 7 dpi, reversing the IL-12/IL-10 ratio during the resolution phase, a potential driver of cellular infiltration. While IL-22 was not present in this mild disease, production of the tissue protective cytokine IL-22 was increased in the serum 7 dpi by neutralization of TGF-β but not IL-4 (**Fig. 8E**). This data suggests that while various overlapping regulatory cytokines inhibit cellular inflammation in the lungs during peak viremia, potentially both IL-4 and TGF-β also inhibit resolution of inflammation around the vessels, but less so around airways and alveoli in this model of mild COVID-19 disease. For IL-4, the persistent inflammation may be through regulation of the IL-12 to IL-10 ratio, though further study is required to determine causality.

**Figure 8.**
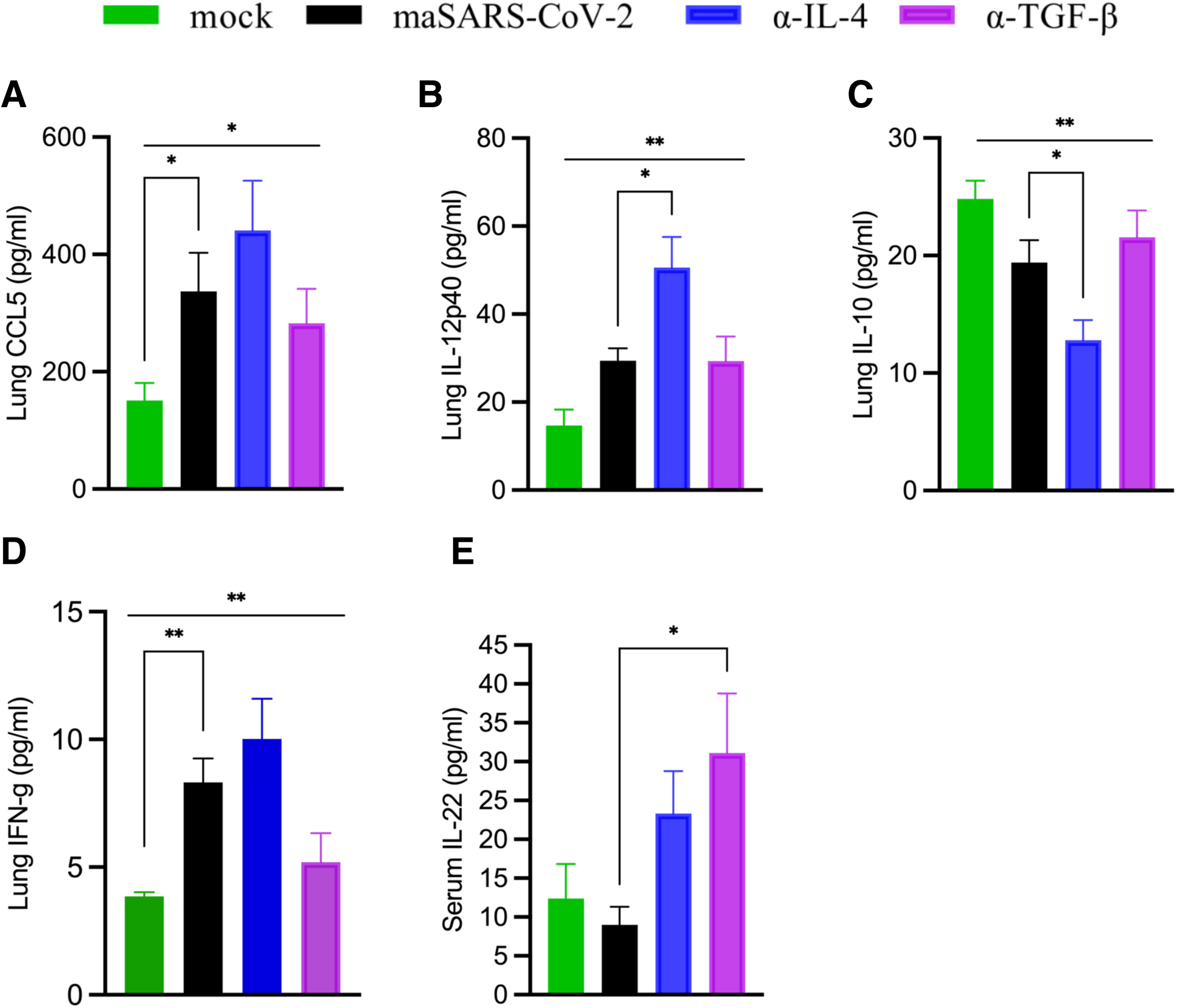
Late Lung and Serum Cytokines. Aged C57BL/6 mice were infected with maSARS-CoV-2, treated with isotype or cytokine neutralizing antibodies on alternate days and weighed daily. On day 7 p.i., the lung homogenate and serum were tested for cytokines by cytokine bead assay or ELISA (IL-33). Graphs show **A)** CCL5; **B)** IL-12p40; **C)** IL-10 and **D)** IFN-γ in lung homogenate, and **E)** serum IL-22 of infected mice. Results are representative of at least two experiments. Statistical analysis was performed by one-way ANOVA followed by Student’s t test. Data were presented as mean ± SEM for groups of five mice. A *p* value at <0.05 was considered statistically significant.

## Discussion

In summary, regulatory cytokines were tested for their ability to affect lung inflammation, weight loss, cytokines and chemokines during infection of aged mice with maSARS-CoV-2. Several studies have been done with this virus and show many similarities with human disease ^28, 44, 45^. Lungs from mice infected with maSARS-CoV-2 showed alveolar thickening and consolidation up to and including pneumonia, as previously published by others in human autopsies and mice ^9, 46–48^. COVID-19 autopsy studies suggest that deceased patients’ lungs fall into three pathological categories which can overlap: ***i.*** acute respiratory distress syndrome characterized by progressive diffuse alveolar damage, ***ii.*** bronchopneumonia resulting from secondary infection with marked neutrophilia, and ***iii.*** tissue thrombosis ^46, 48^. This infection recapitulated the first category displaying excess deposition of extracellular matrix and septal thickening, though we have not quantified fibrosis here. Comparing to Chrabanska et al, hyperplastic type II pneumocytes, hyaline membranes, and thrombosis were present, but variable, similar to their findings ^49^.

Cytokines which correlate with disease severity during human COVID-19 are upregulated in mouse models mirroring quite well those seen in SARS-CoV-2 infection in humans ^50^. We found that maSARS-CoV-2 infected mice had increased production of G-CSF, IL-12 p40, CCL2 and CCL5, which are often linked to monocyte recruitment, a prominent feature of this infection. CCL2 correlated with infiltration of leukocytes seen in lung sections on 3 dpi; while CCL5 and IL-12 p40 also remained elevated on 7 dpi., suggesting a mechanism for prolonged inflammation seen in humans and mice. Vanderheiden *et al* showed a role for inflammatory monocytes in the lungs of animals infected with this same virus. They showed that CCR2 deficiency led to prolonged viremia indicating a role of monocyte-derived cells in viral killing ^28^. They observed large increases in CD86^+^ parenchymal, as distinct from CD86^-^ inflammatory monocyte population. cDC2 (XCR1-CD172a^+^) were also increased in that study, as in human samples. Another study using infection of adeno-associated virus-hACE2 transduced mice also found an expansion of pulmonary infiltrating myeloid-derived inflammatory cells characterized by early Ly6C^hi^ monocytes and inflammatory monocyte-derived macrophages (CD64^+^ CD11c^−^ CD11b^+^ Ly6C^+^) ^44^. While CCL2 is used by monocytes to leave the bone marrow and enter the tissue from the blood, CCL5 likely attracts maturing monocytes to inflamed niches within the tissue ^51^. As IL-12 induces both the chemokines and their receptors, and could promote weight loss, it is a very good candidate for causing the prolonged inflammation seen in the infected mice with IL-4 neutralized, though we have not tested that prediction here ^52^. TGF-β regulates the strength or duration of the immune response, including macrophage activation ^53^, and is essential for lung homeostasis, however its role in COVID-19 has been difficult to ascertain due to low expression levels. TGF-β was reported to increase furin expression leading to enhanced susceptibility to SARS-CoV-2 and disease severity due to increased infection levels, which we did not see here ^54,55^. Most research on TGF-β in lung pathology has focused on fibrosis; however, fibrosis appears to be limited during acute severe COVID-19, and not a likely cause of death from COVID-19. The most common cause of death is respiratory failure due to diffuse alveolar damage, with most patients in the exudative or organizing phase of damage, while later in disease, up to forty percent of patients show fibrosing damage upon autopsy ^56^. To study immune regulation by TGF-β during SARS-CoV-2 infection, we neutralized TGF-β in mice infected with maSARS-CoV-2. Neutralization of TGF-β in mice infected with maSARS-CoV-2 led to the accumulation of Ly6C^hi^ monocytes and decreased the number of CD11c^+^ alveolar macrophages. During infection in the lung, the loss of alveolar macrophages can be compensated by monocyte-derived macrophages potentially explaining the opposite trends ^28, 45^. Furthermore, we detected significantly more IL-6 in the serum of anti-TGF-β-treated mice than isotype-treated group, indicating that IL-6 production is regulated by TGF-β systemically. It would be important to determine the specific steps of monocyte recruitment affected by TGF-β and IL-6, as well as the sources of systemic IL-6 in future studies.

The strongest effects of neutralizing IL-4 was prolonged weight loss and lung cytokines, suggesting a role for IL-4 in the regulation of lung pathology, which was indeed observed. While IL-4-induced type-2 inflammation is rare in viral disease, it is not unprecedented that IL-4 plays a role in viral inflammation for example in RSV infection where RSV G protein preferentially induces a Th2 response contributing to asthma exacerbations ^57^ IL-4 is also known as counter-regulatory to type-1 inflammation ^58^, and can inhibit the synthesis of the proinflammatory mediators TNF-α, IL-1, IL-6, IL-12, PGE_2_, and the chemokines IL-8 and MIP-1 ^35^ as well as the differentiation of type-1 T helper lymphocytes ^36^. In our experiments, neutralization of IL-4 led to increased acute lung inflammation, as evidenced by cellular infiltration and consolidation in the airways of treated mice. Neutralization of IL-4 specifically increased proinflammatory cytokine IL-12p40, and reduced regulatory cytokine IL-10, which combination could promote prolonged inflammation in the lung of maSARS-CoV-2 infected mice. Since IFN-γ was expressed but not significantly affected by neutralization of regulatory cytokines at either timepoint tested, perhaps the effect is more likely to be on monocytic cells than Th1 differentiation. While IL-4 deficient mice have been shown to have an impairment in controlling influenza and other infections due to a reduction in CD8 T cell involvement ^59^, the virus level was not changed in our studies, supporting a primarily immunoregulatory role of IL-4 here. Supporting our conclusions, patients with asthma or helminth infection, type-2 inflammatory diseases of the lung, overall, are less likely to have severe disease and eosinophilia associated with less severe disease ^60^. Eosinophilia, with or without lymphopenia ^31^, is also a common finding in patients admitted for COVID-19, and correlates with severity, but not mortality.

IL-13, a type-2 cytokine that shares the same IL-4 receptor alpha (IL-4Rα) chain with IL-4, has been the subject of several studies in mice and human SARS-CoV-2 infection. IL-13 can be expressed in COVID-19 patients, and levels correlate with the likelihood of requiring a ventilator once hospitalized ^61^. Our data showing that IL-13 is downregulated in this model confirm expression, but do not relate to any role in severity. To study the protective nature of asthma, IL-13 was added to infected human airway epithelial cultures and caused reduced viral infection. IL-5, a driver of eosinophil differentiation, also correlates with a prolonged time to SARS-CoV-2 virus elimination ^62^. An interesting analysis over time in severe versus mild patients in Rwanda suggests that patients who died had, on average more IL-4 and IL-13, as well as IFN-γ, from the earliest timepoints than severely affected patients who lived. Plasma IL-13 and IL-4 are more relevant to severity later in chronic infection, opposite to IL-9, IL-10 and TGF-β, and corresponding with potential counter-regulation of IFN-γ later ^63^. Interestingly, dupilumab, the commercially available anti-IL-4Rα, an antagonist biological drug that blocks both IL-13 and IL-4 activity, reduces disease both in mice, and in patients ^64^. In a small clinical trial, asthma patients taking dupilumab at the time of SARS-CoV-2 infection experienced lower mortality ^61^. Overall, the data suggest a complex interplay of type-2 cytokines in COVID-19 severity, though little is known about IL-4, as distinct from IL-13, or the kinetics of the two cytokines effects in the context of early viremia and longer inflammatory processes. A role for AAM differentiation in immune-pathology has been elegantly verified in SARS-CoV infection using STAT6, STAT1 deficient animals ^30^. While IL-4 and IL-13 have been detected as increased in some mice, in some experiments, particularly on day 2 ^10^, this increase was not observed in this study. This may be related to the extensive use of anesthesia for repeated injections, which did not occur in the other study, but controls to confirm that are lacking. Some mild inflammation was observed in some animals in the mock infected group (not shown).

In this study, we found *Arg1* expression in the lung, suggestive of the presence of alternatively activated monocytic cells. Arg1 can be associated with repair of the lung damage in response to damaging lung parasites like *Nippostrongilus brasiliensis* ^65^, liver fibrosis, and allergic airway inflammation ^66, 67^. As seen here, in other disease states, induction of *Arg1* is dependent upon IL4Rα signaling, and the presence and activation of STAT6. Interestingly, quickly discharged COVID-19 patients have a strong STAT signaling response in peripheral blood mononuclear cells (PBMCs), including pSTAT6 ^68^, the STAT that is specific to IL-4 signaling. This is not surprising, as in SARS-CoV infection, Arg1+ AAM were shown to be important in regulating immune-mediated pathology and prevention of progression to fibrotic lung disease ^69^. The effect of basal IL-4 on Arg1 expression and pathology suggests that additional signals derived from the virus infection, or the immune response to it, are likely required to combine with IL-4 for the effects seen. Similarly, basal levels of IFN-I were shown to be crucial during Listeria infection ^70^. It will also be interesting to determine if AAM are responsible for moderating the recruitment or location of infiltrating cells that are acutely increased in their absence here. It will be important to separate the roles of immunoregulatory cytokines in the earliest phases, on the immune response to virus; and later on, in restoring homeostasis. Overall, our findings suggest multiple novel mechanisms of immunoregulation of the immunopathology underlying the disease process of COVID-19.

## Disclosure

The authors have no financial conflicts of interest.

## Acknowledgements

We are very grateful to the generosity of colleagues and institutions that supported this research during the pandemic including the University of Texas Medical Branch Animal Resources Center staff and veterinarians and the IACUC and biosafety managers and committee members. We thank Dr. Alexander Freiburg for regulatory support, and the excellent discussions of SARS-CoV-2 virology and pathology with the Division of Infectious Diseases, and the Department of Pathology at UTMB, particularly Drs. Susan McClellan and Judith Aronson. We would like to thank Drs. Heidi Risman, Jiayong Xu, John Chan and Padmini Salgame for assistance in BSL3 at Rutgers. We would like to thank Dr. Scott Weaver for facilitating several unique competitive streams of funding.

## Author Contributions

Conceptualization: RS, JJE, SMD, STS, LPC; Investigation: STS, LPC, MGO, TRS, HCT, HGF, MBH, YBMM, KG; Methodology: STS, LPC, MGO, TRS, HCT, HGF, MBH, YBMM, KG; Project Administration: RS, RR, JJE, VDM, MAE; Resources: VDM, MAE, RR, RS, JJE, SMD; Funding Acquisition: RS, RR, JJE, SMD, VDM; Supervision: KG, VDM, MAE, RR, RS, JJE, SMD; Formal Analysis: STS, LPC, MGO, TRS, RR, RS; Visualization: LPC, STS, RS, MJE; Writing – Original Draft Preparation: STS, RS; Writing – Review & Editing: STS, RS, MJE

**Funding** is appreciated from the UTMB Institute for Human Infection and Immunity, the McLaughlin fund, the Sealy Foundation at UTMB, the CII at Rutgers and NIH R01AI135061 (RS), R01AI166668 (RR), 1R01AI153602-01 (VDM) and R21AI145400-A1 (VDM).

## Data Availability

Data supporting the findings of this study are available from the corresponding author, RS, upon request.

## Abbreviations

COVID-19: coronavirus disease 2019
dpi: days post infection
G-CSF: granulocyte-colony stimulating factor
IFN: interferon
maSARS-CoV-2: mouse-adapted SARS-CoV-2
PBS: phosphate-buffered saline
SARS-CoV-2: severe acute respiratory syndrome-coronavirus-2

## Notes

### Competing Interest Statement

The authors have declared no competing interest.

